# Single-Cell Electrophysiological Measurements Reveal Bacterial Membrane Potential Dynamics during Extracellular Electron Transfer

**DOI:** 10.1101/2020.01.15.908145

**Authors:** Sahand Pirbadian, Marko S. Chavez, Mohamed Y. El-Naggar

## Abstract

Extracellular electron transfer (EET) allows microorganisms to gain energy by linking intracellular reactions to external surfaces ranging from natural minerals to the electrodes of bioelectrochemical renewable energy technologies. In the past two decades, electrochemical techniques have been used to investigate EET in a wide range of microbes, with emphasis on dissimilatory metal-reducing bacteria, such as *Shewanella oneidensis* MR-1, as model organisms. However, due to the typically bulk nature of these techniques, they are unable to reveal the subpopulation variation in EET or link the observed electrochemical currents to energy gain by individual cells, thus overlooking the potentially complex spatial patterns of activity in bioelectrochemical systems. Here, to address these limitations, we use the cell membrane potential as a bioenergetic indicator of EET by *S. oneidensis* MR-1 cells. Using a fluorescent membrane potential indicator during *in vivo* single-cell level fluorescence microscopy in a bioelectrochemical reactor, we demonstrate that membrane potential strongly correlates with the electrode potential, produced current, and position of cells relative to the electrodes. The high spatial and temporal resolution of the reported technique can be used to study the single-cell level dynamics of EET not only on electrode surfaces, but also during respiration of other solid-phase electron acceptors.

## Introduction

Respiratory organisms gain free energy by controlling the flow of electrons, from electron donors to electron acceptors through an intricate network of reduction-oxidation (redox) components (1). This electron transport (ET) contributes to the generation of ion motive force (e.g. proton motive force) across a membrane, which is composed of a chemical component (Δ[ion]) and an electrical component (membrane potential, Δψ), and can provide energy for various cellular functions. While molecular oxygen (O_2_) often serves as the terminal electron acceptor for many organisms, many microorganisms are capable of using a variety of other acceptors that can diffuse inside cells to interact with redox components. However, dissimilatory metal-reducing bacteria, such as *Shewanella oneidensis* MR-1, can also transport electrons to insoluble electron acceptors such as metal oxide minerals *outside* the cells (2-6). This extracellular electron transport (EET) process plays an important role in global elemental cycles, and is being harnessed for energy technologies that produce electricity from fuels in microbial fuel cells (MFC), or for production of desirable chemical products from electricity in microbial electrosynthesis (7-9).

EET in *S. oneidensis* is facilitated by an array of multiheme *c*-type cytochromes. Electrons are transferred from the quinone pool to the tetraheme cytochrome CymA at the inner membrane, and ultimately to the MtrABC porin-cytochrome complex that functions as the primary electron conduit across the otherwise insulating cell envelope. This conduit connects the periplasmic decaheme cytochrome MtrA to the outer membrane decaheme cytochrome MtrC through the MtrB outer membrane porin (4, 10-14). MtrC, along with another outer membrane decaheme cytochrome OmcA, act as terminal reductases to transfer electrons to the external electron acceptor, e.g. metal oxide minerals or electrodes. Outside the cell, various mechanisms allow reduction of solid-phase electron acceptors: direct contact with the surface, soluble flavins that can serve either as electron shuttles (15, 16) or as cytochrome-bound redox cofactors enhancing the EET rate (17-19), and via micron-long outer membrane extensions that are proposed to function as bacterial nanowires (20-23). In addition, it was recently shown that the Mtr/Omc cytochromes allow electrons to traverse long distances via a thermally activated redox conduction mechanism along cellular membranes and across multiple neighboring cells (24).

Motivated by the fundamental and technological implications of EET (7, 8), many studies in the past two decades have focused on understanding and enhancing the electron transfer between bacteria and electrodes. These studies have predominantly relied on bulk electrochemical techniques, where typically the goal is to observe and optimize a limited set of outcome variables, e.g. overall current or power output from a whole culture of cells (25). Despite the wealth of information that these techniques provide, their bulk nature precludes an assessment of the inherent complexity within a culture containing billions of cells or more. For instance, these studies typically neglect the likely substantial role of cell-to-cell variability in both EET and cellular energy acquisition. In addition, bulk electrochemical techniques played an important role in investigating the fundamentals of EET mechanisms, including identifying specific proteins involved in EET, through genetic manipulation and experimentation with electrode materials (10, 26). However, it can still be difficult to unambiguously distinguish the mechanism underlying the change in overall electrochemical performance. For example, when a gene encoding a protein of interest is deleted, it can be unclear if the drop in current output is due to the role of that protein in direct electron transfer to electrode, cell attachment, or other factors that play a role in current production.

Addressing existing limitations requires the development of techniques that report EET and cellular activity with simultaneously high spatial (single-cell level) and temporal (dynamic *in vivo* measurements of changing cellular state) resolution during electrochemical measurements. Such a technique could differentiate between the activity of different cells (planktonic vs. attached, varying positions in relation to electrodes) as well as different EET mechanisms (short and long range mechanisms) that contribute to the overall current. This data can then inform the design of bioelectrochemical systems in order to improve their overall performance, or it can help form more direct causal inferences that enhance our understanding of EET mechanisms at a fundamental level.

Although such a technique has been elusive so far, recent studies have made significant progress on improving the spatial resolution in bioelectrochemical techniques. In one group of studies, electrochemical activity of individual cells has been measured using microelectrodes (27-29), revealing the single-cell ET rate and its variability. However, these experiments are prone to low signal-to-noise ratios due to miniscule single-cell currents (∼100 fA), are cumbersome to implement as they involve optical trapping or nanofabrication, and provide no information about the bioenergetic impact on the measured cells. In addition, these measurements generally require removal of cells from their biofilm context and cannot be performed simultaneously on a large number of cells in a bioelectrochemical system. In other experiments, electrodes along with attached cells have been imaged using electron or fluorescence microscopy following electrochemical measurements (10, 16, 26, 30, 31). This end-point imaging, however, does not reveal the *in situ* subpopulation dynamics of EET. In one study, McLean *et al.* (32) addressed these limitations using fluorescence imaging of an optically-accessible microbial fuel cell to estimate the single cell EET by normalizing observed current with the total cellular count on the electrodes. This procedure results in an average cellular EET rate, by assuming no variability in EET or energy gain across the observed population. In other words, it is unable to distinguish between uniform activity by all cells or a much higher level of activity from a small fraction of the population, situations which might necessitate entirely different optimization strategies in bioelectrochemical systems.

Recently, studies on *Bacillus subtilis* biofilms have used *in vivo* measurements of the cell membrane potential to examine the dynamics of metabolic synchronization in biofilms (33-35). A fluorescent cationic dye, Thioflavin T (ThT), was used as a Nernstian membrane potential indicator during live fluorescence microscopy of biofilms without affecting cell viability during days-long experiments. Due to its positive charge, ThT accumulates on the inside of a hyperpolarized (negatively charged) membrane, resulting in an inverse correlation between ThT fluorescence intensity and membrane potential. The high spatial and temporal resolution of this technique demonstrated the role of ionic signaling in *B. subtilis* biofilm synchronization. Motivated by these developments, we set out to test the utility of *in vivo* membrane potential measurements for monitoring the population-wide energetic state under EET conditions on electrodes. Using ThT as a membrane potential indicator during combined *in vivo* fluorescence microscopy and electrochemical measurements, we show that membrane potential can be used as a live indicator of EET activity with single-cell resolution. This technique provides a tool for studying the subpopulation dynamics of EET in microbial electrochemical systems with high spatial and temporal resolution.

## Results

### Thioflavin T as a Membrane Potential Probe in *Shewanella oneidensis* MR-1

Previously, ThT was used to reveal the membrane potential in *Bacillus subtilis* cells (33-35). Similarly, we tested whether ThT fluorescence is a reliable indicator of membrane potential in *S. oneidensis* MR-1. Addition of 10 µM ThT to cells from a late exponential-phase aerobic batch culture resulted in a significant increase in fluorescence intensity of cells (Fig. 1A). To test whether dissipation of membrane potential has an effect on ThT fluorescence, we added the protonophore carbonyl cyanide *m*-chlorophenyl hydrazone (CCCP) to these cells and, as expected, CCCP significantly diminished ThT fluorescence (Fig. 1A). We hypothesized that addition of oxygen to batch-culture cells lacking any electron acceptor would increase the activity of the electron transport chain and in turn hyperpolarize the membrane, leading to an increase in the ThT intensity. Indeed, we observed a significant increase in ThT fluorescence upon addition of oxygen (Fig. 1B). Together, these observations point to ThT as a reliable membrane potential probe in *S. oneidensis*.

**Figure 1:**
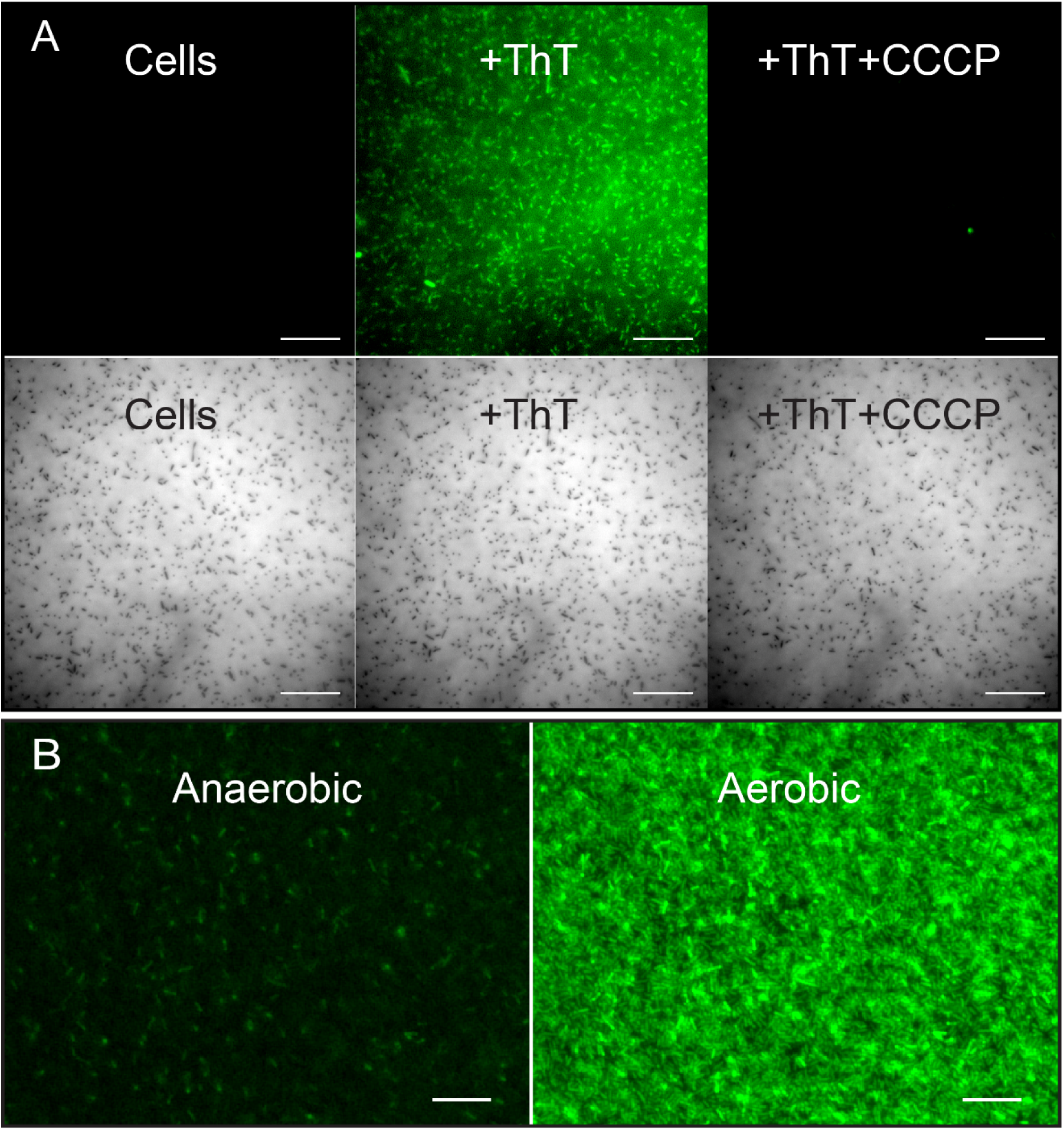
Thioflavin T (ThT) fluorescence is an indicator of membrane potential in *S. oneidensis* MR-1. (**A**) (Left) *S. oneidensis* MR-1 cells shown in fluorescence (top) and brightfield (bottom) channels. (Middle) Addition of ThT to cells significantly enhanced their fluorescence intensity. (Right) Subsequent addition of 125 µM of the protonophore CCCP significantly reduced cell fluorescence. Scale bars are 20 µm. (**B**) Anaerobic *S. oneidensis* culture containing ThT and no electron acceptors (left) shows low cellular fluorescence intensity. Addition of oxygen resulted in a significant increase in fluorescence (right). Scale bars are 10 µm.

### Membrane Potential as an Indicator of Microbe-Anode Electron Transfer

In order to image bacteria inside an electrochemical reactor, we designed a three-electrode bioelectrochemical reactor (Fig. S1) with a working electrode made of indium tin oxide (ITO), an optically transparent, electrically conductive material that has been commonly used in bioelectrochemical systems (17, 24, 27). When placed on an inverted epifluorescence microscope, this reactor allowed for live imaging of cells attached to the electrode surface inside the reactor.

We next used ThT in the bioelectrochemical reactor described above. Anaerobically pre-grown cells were washed in a minimal medium and added to the reactor. The reactor was purged with N_2_ during electrochemical measurements to maintain anaerobic conditions with the electrode serving as the sole electron acceptor for respiration. We repeatedly applied a two-step potential sequence (+0.3 V for 1 hr followed by −0.5 V for 0.5 hr, all potentials vs. Ag/AgCl 1M KCl) to the working electrode while measuring the current and monitoring the ThT fluorescence of cells attached to the electrode. These electrode potentials were chosen to be higher and lower than the redox potentials associated with the EET pathways in *S. oneidensis* MR-1 (18).We consistently observed a strong dependence of ThT fluorescence on electrode potential in wild-type *S. oneidensis* cells, both at the population (Fig. 2A, Movie S1) and single cell (Fig. 2B, Movie S2) levels, with high and low fluorescence tracking the positive and negative potential steps, respectively. Therefore, we hypothesized that the rise in EET activity and produced current during the positive potential step is leading to a more negative membrane potential, in turn causing an increase in ThT fluorescence. To test this hypothesis, we used a mutant, ΔMtr/Δ*mtrB*/Δ*mtrE* (36), lacking genes encoding 8 functional periplasmic and outer membrane cytochromes. As expected, mutant cells produced very little current in the reactor and, consistent with our hypothesis, there was no correlation between electrode potential and ThT fluorescence in mutant cells (Fig. 2C, Movie S1).

**Figure 2:**
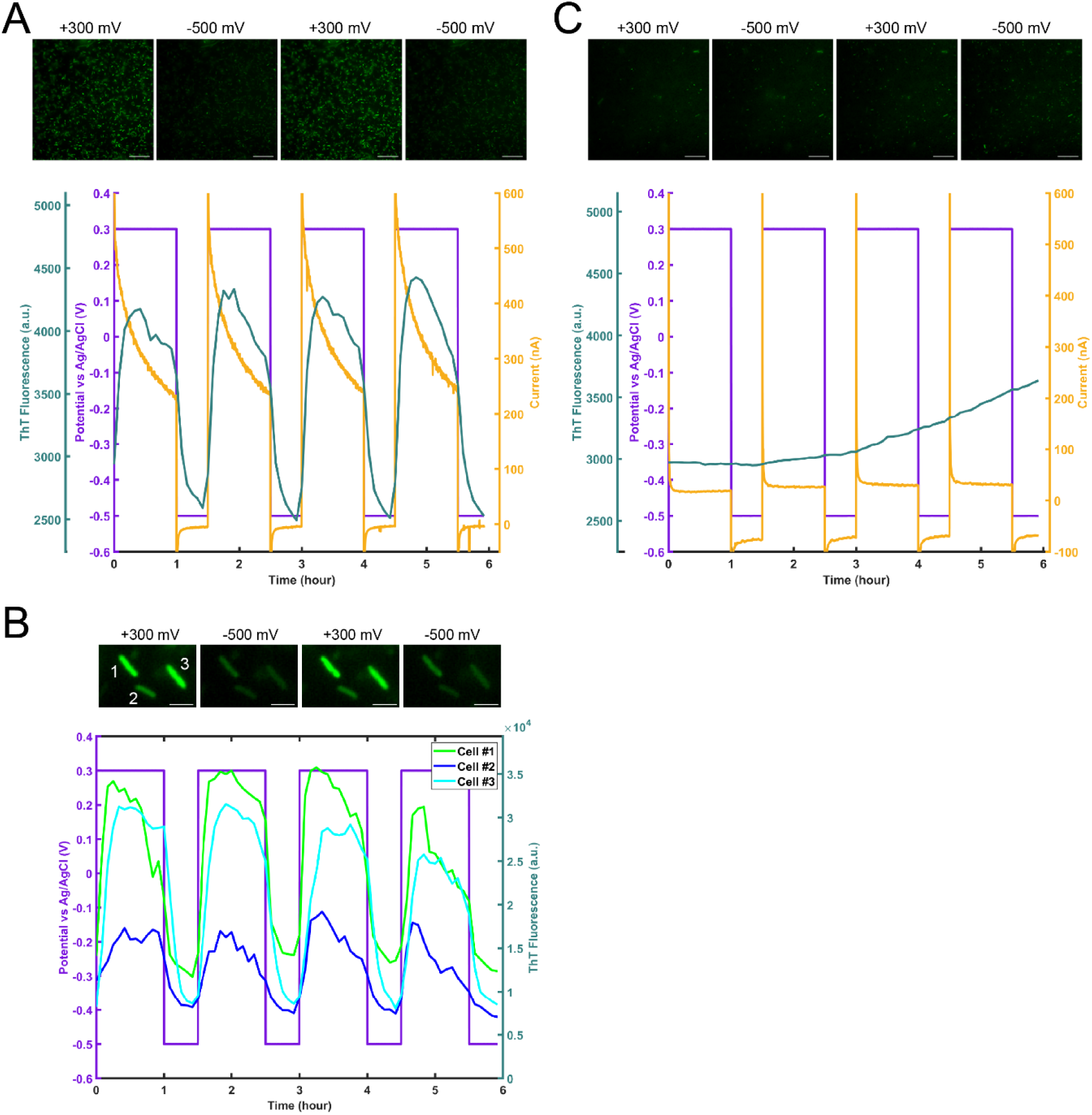
Membrane potential in *S. oneidensis* is an indicator of extracellular electron transfer activity at the single-cell level. (**A**) Fluorescence images along with electrode potential, current, and average Thioflavin T (ThT) fluorescence plots of *S. oneidensis* cells on the ITO working electrode of a bioelectrochemical reactor during a two-step potential sequence (1 hr at +300 mV, 0.5 hr at −500 mV vs Ag/AgCl 1M KCl). Scale bars are 20 µm. (**B**) Fluorescence images along with electrode potential and fluorescence plots of 3 individual *S. oneidensis* cells from (**A**), highlighting the cell-to-cell variability in the larger population. Scale bars are 2 µm. (**C**) Images and plots of the ΔMtr/Δ*mtrB*/ Δ*mtrE* mutant during an identical two-step potential sequence. The mutant strain lacks genes encoding 8 functional periplasmic and outer membrane cytochromes. Scale bars are 20 µm.

To test whether the membrane potential can be monitored continuously in response to a smoothly varying electrode potential, we also performed cyclic voltammetry (CV) in the *S. oneidensis* bioelectrochemical reactors. Similar to the two-step potential sequence described above, ThT fluorescence tracked the cyclic electrode potential (cycle period=26.7 min) with an average 2.9 min ± 1.4 min time lag (Fig. 3, Movie S3).

**Figure 3:**
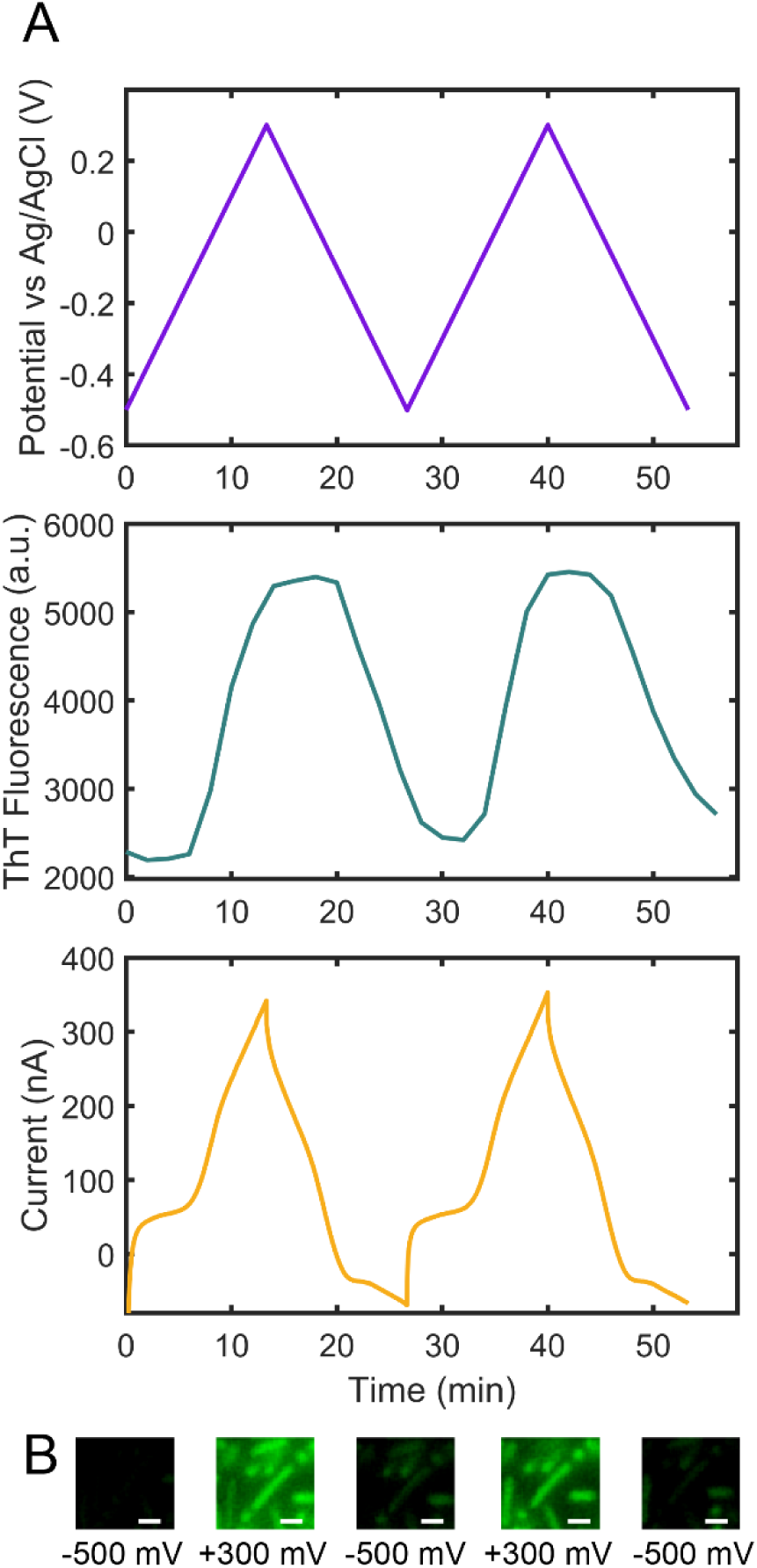
*S. oneidensis* membrane potential is dependent on electrode potential during extracellular electron transfer (EET). (**A**) Electrode potential (vs Ag/AgCl 1M KCl), average cellular Thioflavin T (ThT) fluorescence, and current plots of *S. oneidensis* cells during cyclic voltammetry (CV) in a bioelectrochemical reactor. (**B**) Fluorescence images of individual cells during the CV scans in (**A**), revealing the single-cell level bioenergetic state during bulk CV. Scale bars are 1 µm.

### Membrane Potential as a Function of Microbe-Anode Distance

Next, we examined the correlation between the bioenergetic state (membrane potential probed by ThT) of the cells and proximity to the electrodes. We used ITO patterned chips that provided both a working electrode area (ITO) and an insulating area (glass) in the same field of view. Cells on the electrode showed a strong ThT fluorescence response to changes in electrode potential (Fig. 4, Movie S4), whereas cells on glass did not, except for cells confined within a region near the electrode edge with an average width of 8.9 µm (SD=12.4 µm) (Fig. 4B). To investigate the nature of electrode potential-correlated fluorescence in these cells, we added 5 µM of riboflavin to the reactor during a two-step potential sequence. Flavin addition further expanded the membrane potential region near the electrode by 21.8 ± 4.3 µm (*p*=0.0014) (Fig. 5). We also tested a Δ*bfe* mutant (37), which lacks the bacterial flavin adenine dinucleotide exporter and is inhibited in flavin secretion. The width of the near-electrode region in the Δ*bfe* mutant was not statistically significantly different from the wild type (*p*=0.6690) (Fig. 4B).

**Figure 4:**
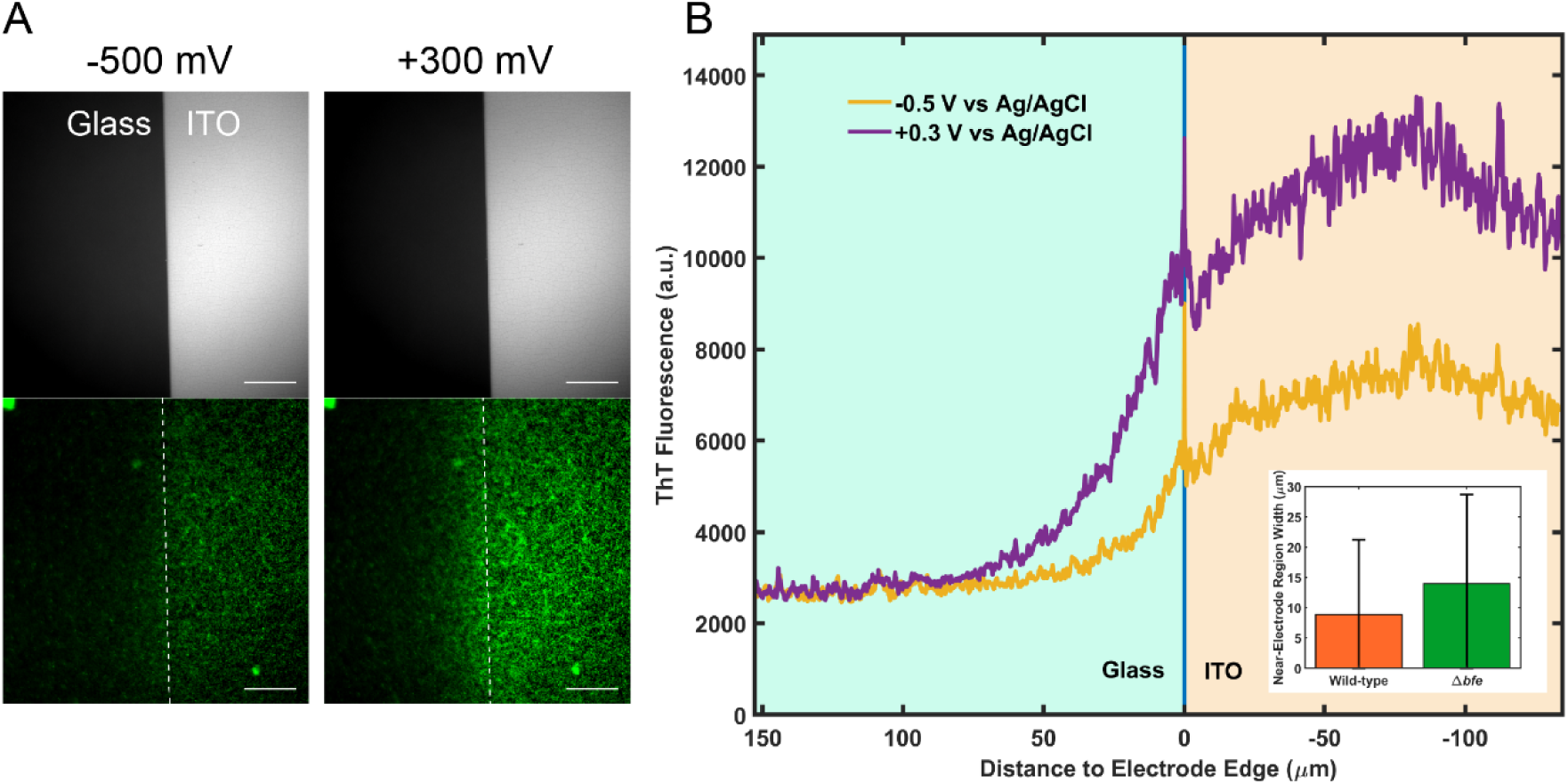
Membrane potential in *S. oneidensis* cells during extracellular electron transfer is a function of microbe-anode distance. (**A**) Brightfield (top) and Thioflavin T (ThT) fluorescence (bottom) images of *S. oneidensis* cells around the boundary between an ITO working electrode (WE) and glass, with the WE at −500 mV (left) and +300 mV (right) vs Ag/AgCl 1M KCl. Scale bars are 50 µm. (**B**) Average ThT fluorescence as a function of distance from ITO-glass edge in fluorescence images of (**A**). (**Inset**) Average width of the near-electrode region of hyperpolarization in *S. oneidensis* MR-1 (wild-type) and a flavin adenine dinucleotide exporter mutant (Δ*bfe*). Error bars represent standard deviation (n=3).

**Figure 5:**
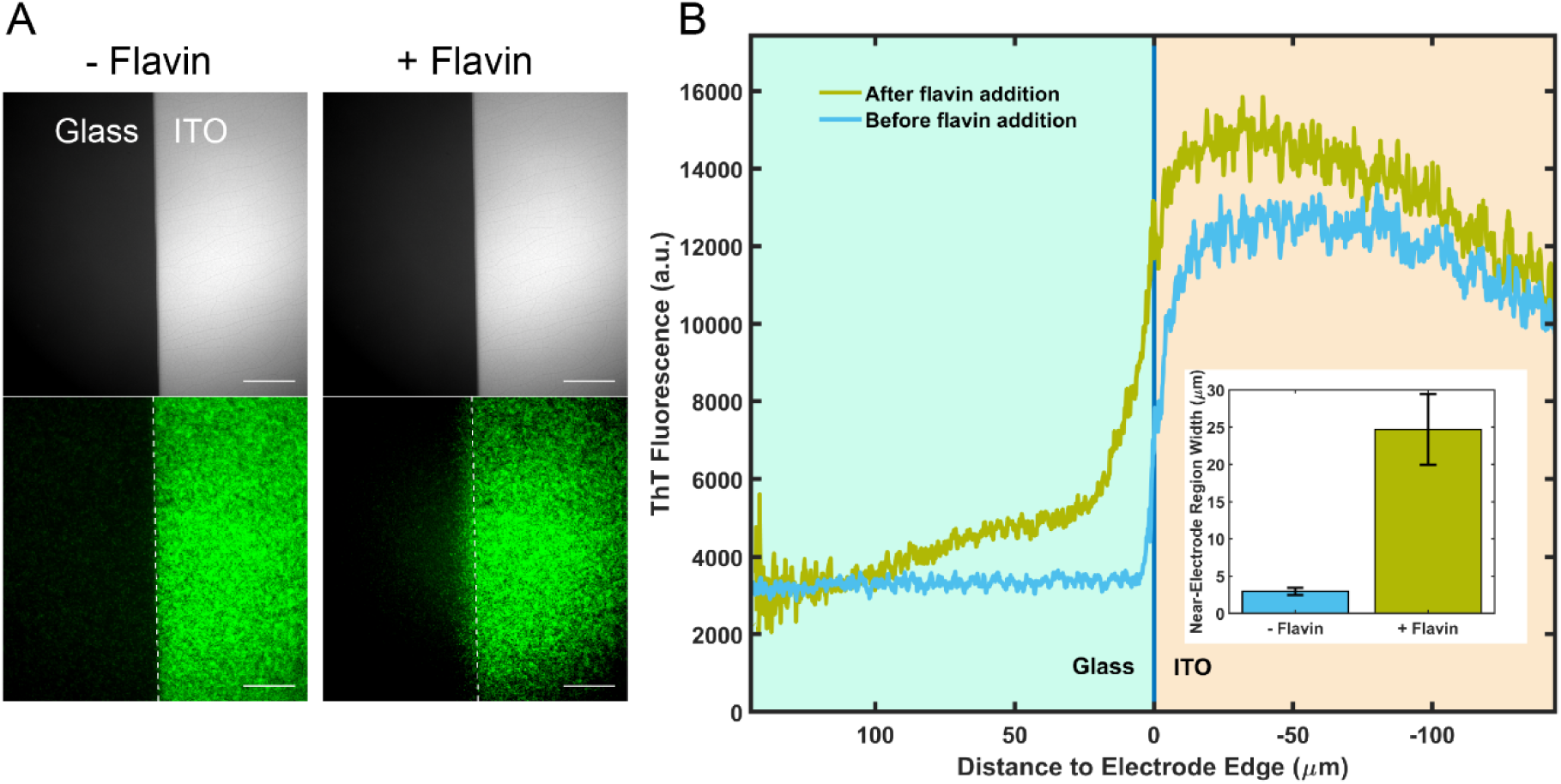
Addition of exogenous riboflavin expands the near-electrode region of hyperpolarization in *S. oneidensis*. (**A**) Brightfield (top) and Thioflavin T (ThT) fluorescence (bottom) images of *S. oneidensis* cells around the boundary between an ITO working electrode (at +300 mV vs Ag/AgCl 1M KCl) and glass, before (left) and after (right) the addition of 5 µM of riboflavin. Scale bars are 50 µm. (**B**) Average ThT fluorescence as a function of distance from ITO-glass edge in fluorescence images of (**A**). (**Inset**) Average width of the near-electrode region of hyperpolarization in *S. oneidensis* before and after the addition of 5 µM of riboflavin. Error bars represent standard deviation (n=3 separate fields of view).

### Effect of Electrode Material on Cellular Activity

The ability to measure EET activity and monitor cellular bioenergetics *in vivo* and at single-cell level presents new opportunities for studying the biotic-abiotic interaction between bacteria and various electrode materials. Gold has been occasionally used as anode material in bioelectrochemial systems (38, 39). However, its efficacy in facilitating EET activity in *S. oneidensis* has been questioned, as it was shown that *S. oneidensis* does not attach well to gold and produces small currents on gold electrodes (26). In order to further assess the suitability of gold, we used patterned gold-coated glass coverslips as working electrodes in our bioelectrochemical reactor. The gold layer was thin enough (5 nm thickness) to allow fluorescence microscopy of cells inside the reactor. Again, we repeatedly applied a two-step potential sequence to the gold electrode and observed that ThT fluorescence of cells on gold, but not on glass, depends on electrode potential (Fig. 6, Movie S5). We also observed cell attachment and currents on gold that were comparable to ITO electrodes (Fig. 6, Movie S5). We therefore concluded that, under our experimental conditions, *S. oneidensis* can efficiently perform EET on gold electrodes.

**Figure 6:**
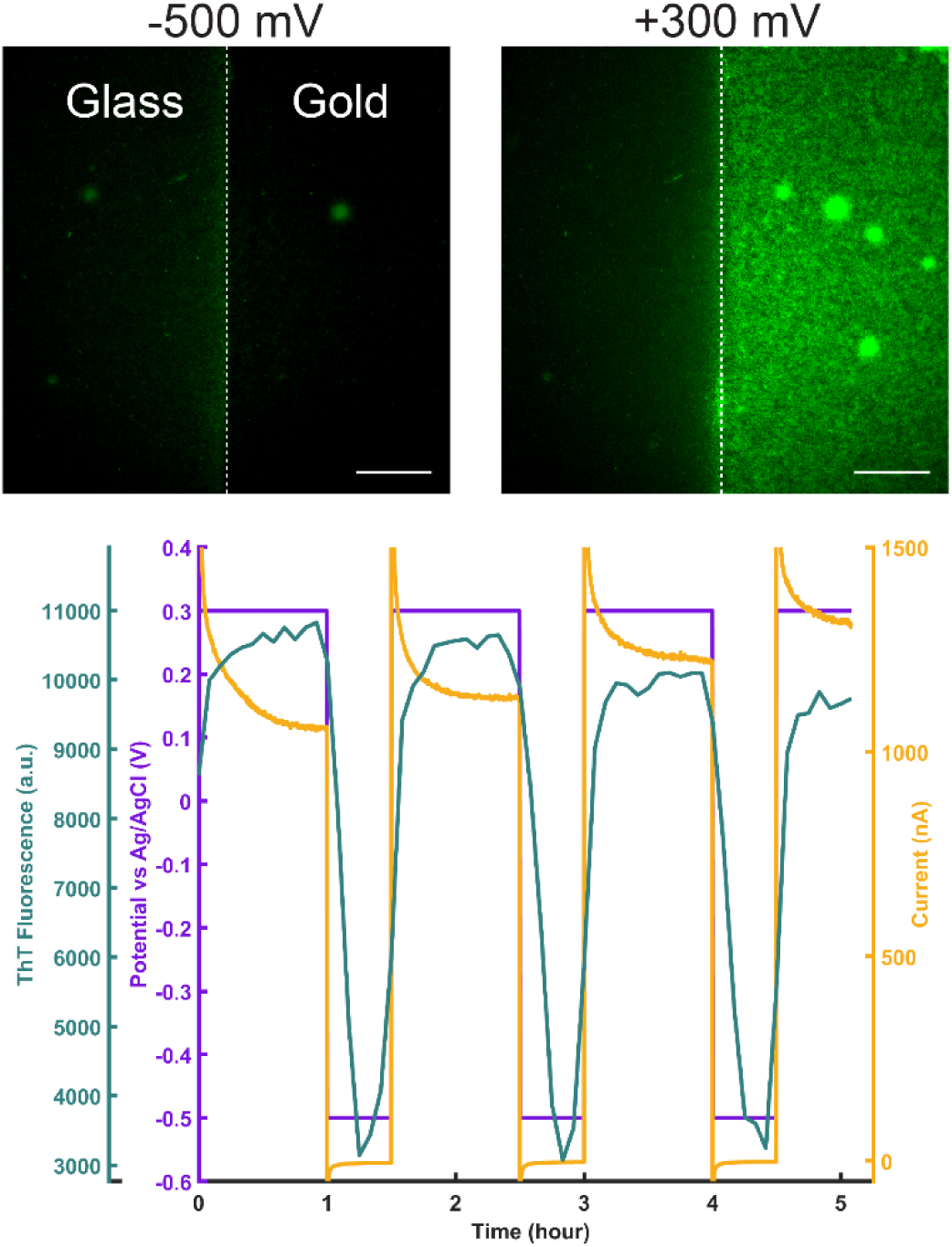
*S. oneidensis* can efficiently perform extracellular electron transfer on gold electrodes. Top: Thioflavin T (ThT) fluorescence images of *S. oneidensis* cells around the boundary between a gold working electrode (WE, right half of each image) and glass (left half of each image), with the WE at −500 mV (left image) and +300 mV (right image) vs Ag/AgCl 1M KCl. Scale bars are 50 µm. Bottom: Electrode potential, current, and average ThT fluorescence plots of *S. oneidensis* cells on a gold working electrode during a two-step potential sequence (1 hr at +300 mV, 0.5 hr at −500 mV vs Ag/AgCl 1M KCl).

## Discussion

We reported a technique for simultaneously measuring microbial EET and tracking the bioenergetic (membrane potential) state at the single cell level. We first showed, by demonstrating the effect of protonophore (CCCP) addition and O_2_ addition, that the fluorescent molecule ThT, previously used to visualize ionic signaling in biofilms, acts as a membrane potential indicator in *S. oneidensis* (Fig. 1). We then applied a periodic two-step potential sequence to the working electrode of a fluorescence microscope-mounted bioelectrochemical reactor and used ThT to show that cellular membrane potential was strongly dependent on electrode potential in the wild-type strain, but not in a mutant lacking the multiheme cytochromes necessary for EET in *S. oneidensis* (Fig. 2, Movies S1 and S2). Additionally, during cylic voltammetry, ThT fluorescence followed the electrode potential with a short time lag relative to the voltammetry cycle period (Fig. 3, Movie S3). This delay can be attributed to the time it takes ThT molecules to traverse the membrane following a change in the membrane potential, and is within the response time range of similar slow-response membrane potential probes (40). Taken collectively, this data demonstrates that membrane potential can be used as an indicator of EET activity in *S. oneidensis*.

The dependence of membrane potential on EET can be understood by noting that, under anaerobic conditions, the *Shewanella* inner membrane is hyperpolarized by proton translocation resulting from the redox cycling of the quinone pool, where quinones are reduced by formate dehydrogenase and lactate dehydrogenase, and quinols are subsequently oxidized by CymA (41). In addition, it was recently shown that while utilizing high potential electron acceptors, *Shewanella* uses proton- and sodium-pumping NADH dehydrogenases that hyperpolarize the membrane and reduce the quinone pool (42). In the presence of an extracellular electron acceptor, electrons are transported from CymA through the Mtr pathway to the acceptor outside the cell, linking the EET at the outer membrane to cation-pumping and thus the membrane potential across the inner membrane. The membrane hyperpolarization resulting from EET serves as an important component of the proton motive force and the sodium motive force (42, 43), providing the free energy for several cellular functions.

The technique described here can be used to investigate the activity of individual cells and help distinguish between different EET mechanisms that factor into overall current production in bioelectrochemical systems. To demonstrate this, we performed the above experiment, combining electrochemical measurements and ThT fluorescence, with patterned working electrodes. The spatial pattern of cellular activity matched well with the electrode pattern, showing that, for the most part, only cells on the electrode contribute to overall current resulting in enhanced membrane potential, and ruling out electron transport beyond ∼30 µm under our experimental conditions (Fig. 4, Movie S4). However, we also observed membrane hyperpolarization in a < 30 μm region near the electrode (Fig. 4). Given the proposed role of flavins as extracellular shuttles and/or cytochrome-bound cofactors that enhance EET in *S. oneidensis*, we suspected that flavins might play a role. When comparing a flavin adenine dinucleotide exporter mutant (Δ*bfe*) with the wild type, we were unable to find a statistically significant difference in the width of the near-electrode region due to the large variability in the size of this region in both strains (Fig. 4B). However, addition of exogenous riboflavin to the wild-type reactor led to a statistically significant expansion of the near-electrode region of hyperpolarization (Fig. 5). This expansion of activity is consistent with more than one hypothesis: (1) Flavins act as electron shuttles between cells on and off electrode, allowing cells in near-electrode region to maintain EET activity; (2) Cells off the electrode transport electrons to electrode via redox conduction through the monolayer of cells as demonstrated previously (24), where added flavins may have enhanced the EET rate by serving as redox cofactors bound to outer membrane cytochromes (18, 19); and (3) Enhanced ionic influx/efflux resulting from higher EET activity of cells on electrode, due to added flavins, impacts the membrane potential of their neighboring cells immediately off the electrode. A similar effect has been observed in *Bacillus subtilis* biofilms, where cells regulate the membrane potential of their neighboring cells by releasing cations (33). To distinguish between these proposals, further investigations are needed to shed light on the mechanism of hyperpolarization in the near-electrode region.

This technique can also directly demonstrate the efficiency of bacterial EET on various electrode materials. For example, we tested the suitability of gold as the anode material in a *S. oneidensis* bioelectrochemical reactor (Fig. 6, Movie S5). Although gold could serve as an ideal anode material for bioelectrochemical studies due to its stability, high conductivity, and ease of fabrication, previous studies on *S. oneidensis* had shown poor attachment and low produced currents on gold electrodes (26, 38, 39). This observation has been attributed to a possible gold toxicity in *Shewanella*. With our technique, we observed that cells can attach and perform EET on gold, evidenced by live measurements of membrane potential and produced current (Fig. 6, Movie S5). Differences in attachment ability and observed EET on gold across studies may therefore reflect different surface properties resulting from the deposition and preparation techniques (Materials and Methods), rather than an intrinsic property of gold itself.

As with any technique, it is important to discuss limitations. While live monitoring of membrane potential can be a powerful tool in studying EET, membrane potential cannot be used as an exclusive indicator of EET activity under all conditions. For example, in the absence of EET, addition of O_2_ to *S. oneidensis* bioreactors results in membrane hyperpolarization (Fig. 1B). Therefore, it is critical that cells are not exposed to O_2_ or other electron acceptors while using membrane potential to study the interaction between cells and a specific acceptor (e.g. an electrode). In addition, ThT fluorescence can be influenced by factors other than membrane potential, such as RNA content (44). Therefore, when using ThT as a membrane potential probe during EET, it is important to control for other contributors to ThT fluorescence, for example, by intentionally manipulating electrode potential as shown in this work.

In summary, we demonstrated dynamic measurements of the bioenergetic state of an electrode-attached population of cells performing EET, with single-cell resolution. By simultaneously performing electrochemical measurements and tracking the membrane potential of the model EET organism *Shewanella oneidensis*, we showed that membrane potential strongly correlates with the electrode potential, EET, and the position of cells relative to the electrodes. Our study opens the possibility that similar techniques may prove useful for studying organisms where the impact of EET on bioenergetics (specifically membrane potential) has not been directly demonstrated, such as newly isolated electrochemically active microorganisms as well as organisms genetically engineered to perform EET.

## Materials and Methods

### Cell Growth and Bioelectrochemical Reactor Inoculation

*Shewanella oneidensis* MR-1, ΔMtr/Δ*mtrB*/Δ*mtrE*, or Δ*bfe* cells were grown from a frozen stock in aerobic LB broth at 30°C, 200 rpm, up to an OD_600_ of 2.1-2.6. 15 mL of the culture was then centrifuged for 5 min at 4226 ×g and resuspended in 10 mL of a defined medium consisting of 50 mM PIPES buffer, 85 mM NaOH, 28 mM NH_4_Cl, 1.34 mM KCl, 4.35 mM NaH_2_PO_4_, 20 mM sodium DL-lactate. The defined medium was supplemented with minerals and amino acids as described previously (10). 2 mL of the resuspended culture was injected into an anaerobic bottle containing 100 mL of the defined medium as well as 20 mM or 40 mM of sodium fumarate. The inoculated bottle was incubated for 20-24 hours at 30°C, 200 rpm. The culture was then centrifuged for 8 min at 7,000 ×g, resuspended in 8 mL of the defined medium, and added to the bioelectrochemical reactor, where cells were allowed to attach to the electrode/glass surface for 20-30 min. The culture in the reactor was then removed (except for the last ∼500 µL which was maintained to avoid drying the attached cells), and replaced by 8 mL of the defined medium that was supplemented by 10 µM of Thioflavin T (ThT).

### CCCP Test

1 mL of the resuspended LB culture (described above) was deposited on a glass coverslip and placed on a Nikon Eclipse Ti-E inverted fluorescent microscope. Cells were imaged before and after the addition of 10 µM of ThT. Subsequently, cells were imaged immediately before and after the addition of 125 µM of the protonophore carbonyl cyanide *m*-chlorophenyl hydrazone (CCCP).

### Bioelectrochemical Reactor Setup

The reactor was made of a 20-mm diameter glass tube glued to the working electrode (planar or patterned ITO or gold on a glass coverslip, as described below) (Fig. S1) using waterproof silicone glue (General Electric Company). The tube was sealed at the top by a custom-made cap that held a Ag/AgCl 1M KCl reference electrode (CHI111, CH Instruments Inc.) and a platinum counter electrode (CHI115, CH Instruments Inc.), as well as the N_2_ inlet and outlet ports (Fig. S1). All electrochemical measurements were performed using a WaveDriver 20 Bipotentiostat/Galvanostat (Pine Research Instrumentation, model# AFP2).

### Electrode Fabrication

The electrodes used in the present study were either purchased from a commercial supplier, fabricated in-house, or designed in-house and then fabricated by a local cleanroom foundry service. Planar ITO-coated glass coverslips were purchased from SPI Supplies (Catalog #:06494-AB). For the in-house fabrication, glass coverslips (24×60 NO. 1 VWR Micro Cover Glasses, Radnor, PA, USA and 43×50 NO. 1 Thermo Scientific Gold Seal Cover Glass, Portsmouth NH, USA) were either rinsed with or sonicated in acetone, isopropanol, and in deionized (DI) water, consecutively. When sonicated, the coverslips sat in each solvent bath for five minutes. The coverslips were then dried with N_2_ and baked on a hotplate at 150°C for ten minutes to remove any remaining water. The coverslips were then placed in a Tegal Plasmaline 515 Photoresist Asher and exposed to an O_2_ plasma at 200 W for 2 minutes.

For the ITO-patterned electrodes, AZ 5214 photoresist (PR) was spin coated and then baked onto the cleaned coverslips. Windows for the ITO pattern were opened in the PR coated coverslips using a Karl Suss MA6 Contact Aligner and a soda-lime glass photomask. A Denton Discovery 550 Sputter Coater was then used to deposit 300 nm of ITO onto the PR coated coverslips. The liftoff of excess ITO was achieved by sonicating the coverslips in one to three consecutive acetone baths for two to ten minutes per bath. The ITO-patterned coverslips were then baked in a N_2_ furnace at 400°C to improve conductivity. For the Au-patterned electrodes, a CHA Industries Mark 40 e-beam and thermal evaporator was used to deposit a 5 nm Ti adhesion layer and then a 5 nm Au layer onto glass coverslips. The Ti and Au layers were both deposited at a rate of 0.02 nm/sec. During the Ti/Au deposition, an approximately 1 cm wide strip of vacuum chamber safe tape was laid across the coverslips. After the deposition, the tape was peeled off, leaving a nonconductive gap between two conductive pads. After fabrication, the coverslips were then rinsed with acetone, isopropanol, DI water, dried with N_2_, and baked on a hotplate at 150°C for ten minutes. The coverslips were then exposed to an O_2_ plasma at 100 W for 1 minute. Additional ITO-patterned electrodes were fabricated using standard cleanroom photolithography, similar to the above description, at the University of California, San Diego Nano3 cleanroom.

### Simultaneous Electrochemical and Fluorescence Measurements

The bioelectrochemical reactor containing the culture was placed over a 40× or a 100× objective of a Nikon Eclipse Ti-E inverted fluorescent microscope equipped with a drift correction unit (Nikon Perfect Focus System). N_2_ flow into the reactor was started just before the beginning of the measurements. Time-lapse microscopy and electrochemical measurements were started simultaneously. In the two-step potential sequence measurements, +0.3 V for 1 hour followed by −0.5 V for 0.5 hour (vs Ag/AgCl, 1 M KCl) was repeatedly applied on the working electrode, while brightfield and ‘FITC’ (Nikon filter set B-2E/C) fluorescence images were acquired at 5-min intervals. In cyclic voltammetry measurements, the working electrode potential was swept at 1 mV/s, while brightfield and ‘FITC’ images were acquired at 2-min intervals. In the riboflavin addition experiment, 5 µM riboflavin was added to the bioelectrochemial reactor containing *S. oneidensis* MR-1 cells during a two-step potential sequence and simultaneous fluorescence microscopy.

### Oxygen Addition Test

The experimental setup was similarly prepared as in the electrochemical measurements, but without any potential applied on the working electrode. Fluorescence imaging began with N_2_ flow into the reactor maintaining anaerobic conditions. After 1.5 hour, the N_2_ flow was stopped and was replaced by ambient air flow. Fluorescence images were acquired immediately before and after the change from N_2_ to air.

### Image Analysis

Fluorescence images acquired by the microscope software (NIS-Elements AR 4.60, Nikon Inc.) were exported as 16-bit images after linear brightness and contrast adjustments were made to each entire image, with identical adjustments in all images from the same time-lapse experiment. Images were then imported into MATLAB (R2019a, Mathworks). A custom MATLAB code was used to extract an average cell fluorescence intensity in each image by subtracting the background fluorescence, excluding pixels with intensities lower than a set threshold in order to retain only cell fluorescence, and averaging the intensity of the remaining pixels. In the riboflavin addition experiment, different brightness and contrast adjustments were made to images from before and after riboflavin addition due to the large increase in fluorescence background upon riboflavin addition. Therefore, the fluorescence intensities in these images were not directly compared with each other. Instead, these images were only used independently to calculate the width of the near-electrode region, which depends on the relative fluorescence intensities of pixels on glass versus electrode within the same image.

To quantify the average pixel intensity as a function of distance from the electrode, the electrode-glass edge was manually defined based on the corresponding brightfield image, in turn allowing for the calculation of the distance from each pixel to the edge. The average pixel intensity was then plotted as a function of distance, with negative and positive distances indicating pixels on the electrode and glass, respectively.

### Width of the Near-Electrode Region of Hyperpolarization

The near-electrode region width (*w*) was defined as the distance from the electrode-glass edge where the fluorescence intensity on glass drops by a factor of *e* (∼2.718) relative to the electrode fluorescence intensity. This can be written as *fl*(*w*)=(*fl*_*electrode*_-*fl*_*glass*_)/*e*, where *fl*(*w*) is the fluorescence intensity on glass at the end of the near-electrode region, *fl*_*electrode*_ is the average fluorescence intensity on the electrode (or the fluorescence intensity at the electrode edge in the riboflavin addition experiment), and *fl*_*glass*_ is the average fluorescence intensity on glass far from the electrode (50 µm and farther from the electrode edge).

### Statistical Analysis

For the comparison of the near-electrode region width between *S. oneidensis* MR-1 and Δ*bfe* (n=3 separate experiments for each strain), or between before and after riboflavin addition (n=3 separate fields of view from the same experiment), two-sided two-sample Student’s *t*-test was performed to evaluate statistical significance, at a significance level of 0.05.

## Acknowledgments

We thank Dr. Jeffrey Gralnick for kindly providing the ΔMtr/Δ*mtrB*/Δ*mtrE* mutant strain. We also thank the Nanoelectronics Research Facility at the University of California Los Angeles and the Nano3 Cleanroom at the University of California San Diego for making the fabrication of the electrodes possible. This study was supported by the U.S. Office of Naval Research Multidisciplinary University Research Initiative Grant No. N00014-18-1-2632. We also acknowledge support for the development of the technique from the Air Force Office of Scientific Research Presidential Early Career Award for Scientists and Engineers (FA955014-1-0294, to M.Y.E.-N.).

## Supplementary Materials

**Fig. S1.**
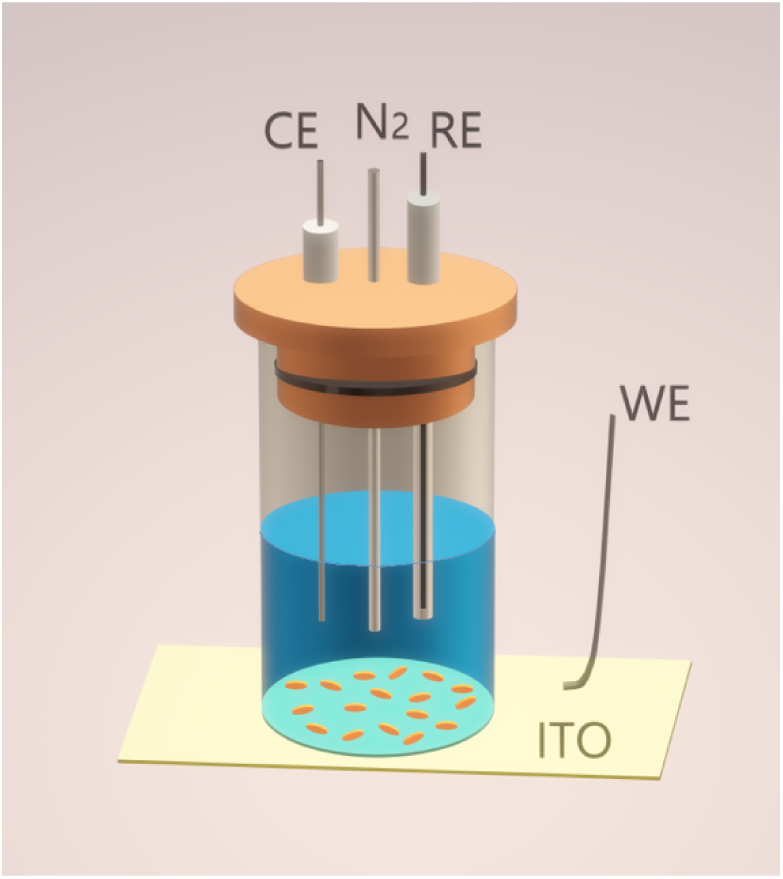
Schematic of the bioelectrochemical reactor. The reactor consisted of a glass tube attached to a working electrode (WE, ITO or gold-coated glass coverslip) at the bottom and sealed with a cap holding the counter electrode (CE), the reference electrode (RE), and the N_2_ port. Cells attached to the transparent WE were imaged from below by an inverted fluorescent microscope.

## Captions for Supplementary Movies

Movie S1: **Thioflavin T (ThT) fluorescence time-lapse movie of *S. oneidensis* wild-type and *ΔMtr/ΔmtrB/ΔmtrE* mutant cells on an ITO working electrode of a bioelectrochemical reactor**. (**Left**) Fluorescence time-lapse movie along with the electrode potential, current, and average ThT fluorescence plots of *S. oneidensis* cells on the working electrode during a two-step potential sequence (1 hr at +300 mV, 0.5 hr at - 500 mV vs Ag/AgCl 1M KCl). (**Right**) Movie and plots of the ΔMtr/Δ*mtrB*/Δ*mtrE* mutant (‘cytochrome mutant’) during an identical two-step potential sequence. The mutant strain lacks genes encoding 8 functional periplasmic and outer membrane cytochromes. Scale bars are 20 µm.

Movie S2: **Thioflavin T (ThT) fluorescence time-lapse movie of individual *S. oneidensis* cells on an ITO working electrode of a bioelectrochemical reactor**. Fluorescence time-lapse movie along with electrode potential and fluorescence plots of 3 individual *S. oneidensis* cells on the working electrode during a two-step potential sequence (1 hr at +300 mV, 0.5 hr at −500 mV vs Ag/AgCl 1M KCl). Scale bar is 2 µm.

Movie S3: **Thioflavin T (ThT) fluorescence time-lapse movie of *S. oneidensis* cells during cyclic voltammetry (CV) in a bioelectrochemical reactor**. Fluorescence time-lapse movie along with electrode potential (vs Ag/AgCl 1M KCl), current, and average ThT fluorescence plots of *S. oneidensis* cells during CV in a bioelectrochemical reactor. The CV scan rate was 1 mV/s. Scale bar is 20 µm.

Movie S4: **Thioflavin T (ThT) fluorescence time-lapse movie of *S. oneidensis* cells on a patterned-ITO working electrode of a bioelectrochemical reactor**. The movie shows cells around the boundary between the ITO working electrode (right half) and glass (left half) during a two-step potential sequence (1 hr at +300 mV, 0.5 hr at −500 mV vs Ag/AgCl 1M KCl). The plots show the average ThT fluorescence as a function of distance from the ITO-glass edge in the movie. The yellow line on the plots indicates the ITO-glass edge. The working electrode potential (vs Ag/AgCl 1M KCl) in each frame is shown at the top. Scale bar is 50 µm.

Movie S5: **Thioflavin T (ThT) fluorescence time-lapse movie of *S. oneidensis* cells on a patterned-gold working electrode of a bioelectrochemical reactor**. The movie shows cells around the boundary between the gold working electrode (right half) and glass (left half) during a two-step potential sequence (1 hr at +300 mV, 0.5 hr at −500 mV vs Ag/AgCl 1M KCl). The plots show the average ThT fluorescence as a function of distance from the gold-glass edge in the movie. The yellow line on the plots indicates the gold-glass edge. The working electrode potential (vs Ag/AgCl 1M KCl) in each frame is shown at the top. Scale bar is 50 µm.

